# Combination of ipratropium bromide and salbutamol in children and adolescents with asthma: a meta-analysis

**DOI:** 10.1101/2020.07.31.230318

**Authors:** Hongzhen Xu, Lin Tong, Peng Gao, Yan Hu, Huijuan Wang, Zhimin Chen, Luo Fang

**Author notes:** **Correspondence authors** *Luo Fang, PhD* Department of Pharmacy, The Children’s Hospital, Zhejiang University School of Medicine, National Clinical Research Center for Child Health, Hangzhou, Zhejiang, China *Zhimin Chen, MD, PhD* Department of Pulmonology, The Children’s Hospital, Zhejiang University School of Medicine, National Clinical Research Center for Child Health, Hangzhou, Zhejiang, China.

## Abstract

**Background:** A combination of ipratropium bromide (IB) and salbutamol is commonly used to treat asthma in children and adolescents; however, there has been a lack of consistency in its usage in clinical practice.

**Objective:** To evaluate the efficacy and safety of IB + salbutamol in the treatment of asthma in children and adolescents.

**Methods:** The MEDLINE, Embase, and Cochrane Library as well as other Chinese biomedical databases (including China Biological Medicine Database, Chinese National Knowledge Infrastructure, Chongqing VIP, and Wanfang Chinese language bibliographic database) were systematically searched from the date of database inception to 02/09/2019 for randomized controlled trials in children and adolescents (≤18 years) with asthma who received IB + salbutamol or salbutamol alone. The primary outcomes included hospital admission and adverse events. A random effects model with a 95% confidence interval (CI) was used. Subgroup analysis was performed according to age, severity of asthma, and co-interventions with other asthma controllers. This study was registered with PROSPERO.

**Results:** Of the 637 studies that were identified, 55 met the inclusion criteria and involved 6396 participants. IB + salbutamol significantly reduced the risk of hospital admission compared with salbutamol alone (risk ratio [RR] 0.79; 95% CI 0.66–0.95; p = 0.01; I^2^ = 40%). Subgroup analysis only showed significant difference in the risk of hospital admission in participants with severe asthma exacerbation (RR 0.71; 95% CI 0.60–0.85; p = 0.0001; I^2^ = 0%) and moderate-to-severe exacerbation (RR 0.69; 95% CI 0.50–0.96; p = 0.03; I^2^ = 3%). There were no significant differences in the risk of adverse events between IB + salbutamol group and salbutamol alone group (RR 1.77; 95% CI 0.63–4.98).

**Conclusion:** IB + salbutamol may be more effective than salbutamol alone for the treatment of asthma in children and adolescents, especially in those with severe and moderate to severe asthma exacerbation. Future prospective research on these subgroup population are needed.

## Introduction

Asthma is the most common chronic disease among children and is estimated to affect 300 million individuals worldwide ^1^. In China, asthma affects 3% of children ≤14 years of age and the prevalence of childhood asthma has increased by 50% over the past 10 years ^2^. Asthma-related hospitalization can negatively affect the quality of life of children and their caregivers. Additionally, health care expenditures for asthma-related conditions impose considerable economic burden on society ^3; 4^.

Almost all available guidelines recommend that the repeated administration of inhaled short-acting β_2_-agonists (SABAs, up to 4–10 puffs every 20 minutes for the first hour) is an effective and efficient way to achieve rapid reversal of airflow limitation in patients with mild-to-moderate asthma exacerbation ^2; 5^. Currently, several available guidelines ^6–9^ have recommended the addition of ipratropium bromide (IB), a short-acting muscarinic acetylcholine receptor antagonist, to SABAs as an optional treatment for children and adolescents with acute asthma exacerbation. Although IB does not seem to be very efficient in controlling asthma, several studies have demonstrated that a combination of IB and SABAs is associated with fewer hospitalizations and greater improvement in peak expiratory flow (PEF) and forced expiratory volume in one second (FEV_1_) compared with SABA alone in children and adolescents with moderate-to-severe asthma exacerbation ^10–16^. However, these recommendations lack uniformity with respect to the optimal age, severity of asthma, and co-intervention with other asthma controllers for such therapy.

The most recent systematic review comparing IB + SABA and SABA alone for the treatment of acute asthma in children and adolescents was published in 2013 and reported a combined treatment benefit as evidenced by a decrease in the risk of hospital admission and improved lung function and clinical scores ^14^. However, the review found no effect of age and co-intervention (such as steroid or standard care) on the hospital admission rate to treatment.

Since this last publication, there have been numerous studies published, and thus, this systematic review and meta-analysis was performed to update the evidence comparing salbutamol alone with IB + salbutamol for identifying the impact of the combination treatment in children and adolescents with asthma

## Results

### Results of the search

The initial electronic database search identified a total of 632 references. Another five references were identified after checking the references listed in the relevant systematic reviews and included studies. After removing duplicate publications, 607 studies were included. After evaluating the titles and abstracts at first-level screening, 87 records were included. Assessment of the full text at second-level screening removed another 32 records. Finally, 55 RCTs were included. These RCTs involved 6396 participants and met the inclusion criteria for this review (Figure 1) (for full references, refer to E-Appendix 3).

### Characteristics of included studies

The included 55 RCTs (53 published trials and 2 unpublished trials ^19; 20^) were from both developing and developed countries, including Australia, China, Canada, Chile, Greece, India, Mexico, Pakistan, Spain, Turkey, Thailand, the United Kingdom, and the United States. All trials included pediatric patients. The age group varied across studies from 4 months to 18 years. Asthma severity varied from mild to severe on different scales across the trials. Co-interventions were administered in 31 studies, with 17 of them combining glucocorticoid, 3 with steroids, and 11 with standard care (E-Appendix 2). The frequency of IB + salbutamol treatment ranged from every 10 minutes to every 24 hours. Moreover, 23 studies reported that the treatment duration of IB + salbutamol was less than 120 minutes (median = 60 minutes), 18 studies reported that treatment duration ranged between 3 days and 40 weeks (median = 7 days); and 14 studies did not report treatment duration (E-Appendix 4). All characteristic information was collected based on reported data from original studies.

### Risk of bias in the included studies

Quality analysis was performed on the basis of aforementioned methods and tools. Details of the risk of bias assessment are provided in Figure 2. Only one study was assessed as being at low risk of bias in all domains ^10^. Five studies were considered to be at high risk of bias, one of which was due to random sequence generation ^21^, two due to blinding setting ^17; 22^ and the other two due to selective reporting on predefined outcomes ^12; 23^. The remaining 49 studies were considered to be at unclear risk of bias (for details, refer to E-Appendix 5).

### Primary outcome

Hospital admission was reported by 16 trials involving 2834 participants ^10; 11; 13; 19–22; 24–32^. The meta-analysis was conducted with 15 trials showed that compared with salbutamol alone, the IB + salbutamol group showed a significant reduction in the risk of hospital admission (RR 0.79; 95% CI 0.66–0.95; I^2^ = 40%; p = 0.01; Figure 3). One study ^21^ was not included in the meta-analysis because it reported a number of zero on hospital admission in intervention and comparison groups.

Regarding the subgroup analysis, there was a significant difference in hospital admission according to severity of illness (test for subgroup differences: χ^2^ = 14.34, df = 6, p = 0.03, I^2^ = 58.2%). Furthermore, administration of IB + salbutamol only showed a significant reduction in the hospital admission in participants with severe asthma exacerbation (RR 0.71; 95% CI 0.60–0.85; p = 0.0001; I^2^ =0%; 1270 participants in ten studies^10; 11; 24–26; 29; 30–32^) and moderate-to-severe asthma exacerbation (RR 0.69; 95% CI 0.50–0.96; p = 0.03; I^2^ = 3%; 629 participants in four studies ^11; 13; 19; 20^). There were no significant differences between IB + salbutamol and salbutamol alone in participants with mild asthma exacerbation (RR 1.43; 95% CI 0.42–4.79; p = 0.57; 117 participants in one study ^32^), moderate asthma exacerbation (RR 1.04; 95% CI 0.89–1.22; p = 0.59; I^2^ = 2%; 736 participants in four studies ^10; 11; 31; 32^), and mild-to-moderate asthma exacerbation (RR 0.85; 95% CI 0.51–1.43; p = 0.54; 348 participants in two studies ^19; 27^). Additionally, there were no significant differences in the age subgroup (χ^2^ = 1.20, df = 3, p = 0.75, I^2^ = 0%) or the co-intervention subgroup (χ^2^ = 0.88, df = 4, p = 0.93, I^2^ = 0%) (for details, refer to E-Appendix 6).

Eight trials (with 1137 participants) reported the number of participants who had adverse events. ^28; 31; 33–37^. Three trials reported a number of zero on adverse events in both intervention and comparison groups ^33; 37–38^. Based on reporting in the remaining five trials ^28; 31; 34–36^, 65 participants had adverse events in the IB + salbutamol group (with 349 participants), and 47 participants had adverse events in the salbutamol alone group (with 348 participants).The results of meta-analysis on these five trials (with 697 participants) ^28; 31; 34–36^ showed no significant differences on the incidence of adverse events between the compared groups (RR 1.77; 95% CI 0.63–4.98; p = 0.28; I^2^ = 77%, Figure 4). The substantial heterogeneity may be explained by the different treatment durations among the five studies in the meta-analysis. In two of the studies ^28,31^ patients were treated with IB + salbutamol for 60–90 minutes, whereas in the other three studies ^34–36^, patients were treated for 3–7 days. The differences in treatment durations may have led to clinical heterogeneity.

There were no significant differences in the subgroup analysis for the incidence of adverse events between IB + salbutamol and salbutamol alone in the severity subgroups (test for subgroup differences: χ^2^ = 1.49, df = 1, p = 0.22, I^2^ = 32.7%), age subgroups (test for subgroup differences: χ^2^ = 0.88, df = 2, p = 0.65, I^2^ = 0%), and co-intervention subgroups (test for subgroup differences: χ^2^ = 3.23, df = 2, p = 0.20, I^2^ = 38.1%). Only one study ^31^ including 347 participants with moderate asthma exacerbation reported a significant reduction in the number of adverse events in the salbutamol alone group compared with the IB + salbutamol group (RR 2.86; 95% CI 1.31–6.21; p = 0.008) (for details, refer to E-Appendix 7).

### Secondary outcome

Pulmonary function was reported in 5 studies (Table 1), among which three reported predicted % FEV_1_ at both 60 min and 120 min ^19; 28; 30^after treatment, and one reported predicted % FEV_1_ at 60min ^20^ only. Among these four studies, two studies ^28; 30^ significantly favoring IB + salbutamol therapy at 120 min after treatment. One study ^30^ significantly favoring IB + salbutamol therapy at 60 min after treatment. Only one study reported absolute % FEV_1_ at 60 min and 120 min ^40^, favoring IB + salbutamol therapy at 120 min after treatment. Another study reported respiratory resistance at 60 and 120 min ^19^, with no significant difference between IB + salbutamol and salbutamol alone groups.

Nine studies reported clinical scores regarding different symptoms at various timepoints ranging from 15 min to 240 min, with one study did not report treatment duration ^43^ (E-Appendix 8). Among them, one study reported dyspnea scores ^41^; one reported respiratory distress scores ^42^; five reported wheeze scores ^12; 39;40; 41; 43^; two reported asthma scores ^11; 27^; two reported wheezing sound scores ^40; 41^; one reported cough scores ^41^; and one reported clinical scores based on clinical examination, activity, and speech ^44^, with three of them significantly favoring IB + salbutamol therapy ^11; 41; 43^.

Regarding various specific adverse events (E-Appendix 8), dry month^28; 35; 40; 43; 45; 46^, nausea^13; 19; 20; 28; 30; 31^, tremor^20; 22; 29; 31; 43; 45;47^, and vomiting^19; 20; 22; 26; 28; 29; 31; 47^ were reported in more than two trials. IB + salbutamol group showed significant reduction on the incidence of nausea compared with salbutamol alone group (RR 0.60; 95% CI 0.39, 0.93; p=0.02; I^2^=0%; seven studies with 763 participants). However, none of the other three outcomes showed significant differences between the two groups (p > 0.05). In addition, there was also no significant difference in other adverse events (such as abdominal pain, headache, palpitations, etc) between the two groups (p > 0.05) (for details, refer to E-Appendix 8).

Additionally, there was no significant difference in oxygen saturation (p = 0.18), need for extra medication (repeated bronchodilator treatments (p = 0.32), systemic corticosteroids (p = 0.41)), and relapse rate (p = 0.85) between the two groups (E-Appendix 8).

### Sensitivity analysis

In sensitivity analysis omitting enrolled studies in turn, the results remained consistent across different analyses, which suggested that the findings were reliable and robust (for details, refer to E-Appendix 9).

### Publication bias

Publication bias of the studies was assessed using funnel plots for hospital admission and relapse rates. No obvious asymmetry was observed in all groups (E-Appendix 10).

## Discussion

There is a lack of consistency in clinical practice regarding the treatment of asthma exacerbation in children and adolescents. Since the last systematic review published in 2013 ^14^, a considerable number of new studies evaluating the efficacy and safety of IB + salbutamol compared with those of salbutamol alone for the treatment of asthma exacerbation in children and adolescents have been published. This systematic review was conducted to update the findings on this topic and provide clinicians with the most current information to aid in the decision-making process involved in determining the best treatment options for the pediatric population presenting with acute asthma exacerbation.

This systematic review supported the benefits of IB + salbutamol for the treatment of asthma in children and adolescents according to the reduction in hospital admission (RR 0.79; 95% CI 0.66–0.95). We performed subgroup analysis to determine whether age, severity of asthma, and co-intervention influenced the effect of IB + salbutamol treatment on hospital admission. Although the subgroup analyses might have contained overlap and non-randomized participants, the result could probably suggest the benefits in children and adolescents with severe asthma exacerbation (RR 0.71; 95% CI 0.60–0.85) and moderate-to-severe asthma exacerbation (RR 0.69; 95% CI 0.50–0.96), which is consistent with the results of a previous systematic review ^14^. Consistent with the findings of Castro-Rodriguez (2015) ^48^, patient age did not alter the effect of IB + salbutamol on reduction of the risk of hospital admission. However, contrary to the findings of Griffiths (2013) ^14^, IB + salbutamol showed no significant reduction in the risk of hospital admission in patients with co-intervention of glucocorticoid. A possible explanation is the difference in interventions between the present review and Griffiths’ (2013) ^14^ review. The previous review reported a wider range of intervention that included all types of combined inhaled anticholinergics and SABAs, which may have included studies focused on terbutaline. However, the present review only included IB + salbutamol as an intervention treatment. Therefore, studies with a focus on terbutaline were excluded. Another explanation could be the updated search date. Compared with the review by Griffiths (2013) ^14^, the present review included additional 6-year literature published between 2013 and 2019. Moreover, Griffiths (2013) ^14^ used a fixed effects model to analyze data, whereas the present study used a random effects model. The use of different statistical models may also explain the difference in the results.

Consistent with previous systematic reviews^14,48^, IB + salbutamol could significantly improve the predicted % and absolute % change in FEV_1_ at both 60 and 120 minutes after treatment compared with salbutamol alone. Contrary to the review by Griffiths (2013) ^14^, the increase in lung function observed with the combined treatment was not associated with an increase in oxygen saturation. A possible explanation for this is that the pervious review ^14^ used oxygen saturation < 95% instead of percentage of oxygen saturation as the outcome indicator.

Consistent with previous systematic review^14^, nausea, vomiting, and tremors were listed as secondary outcomes because of the direct treatment-related effects of salbutamol or ipratropium bromide. Although the combination of IB and salbutamol was previously found to result in fewer tremors and less nausea compared with salbutamol alone ^16,17^, we found consistent results in nausea (RR 0.60; 95% CI 0.39–0.93) but did not identify any significant difference in the other three adverse events between the two groups. Possible explanations could be what has mentioned previously for subgroup results.

The present review found that most Chinese studies reported a clinical response as an outcome after treatment using IB + salbutamol or salbutamol alone in children and adolescents. However, because the clinical response was not clearly defined in the original studies, it was not included as a secondary outcome in the present review.

A key contribution of this present review is the update of evidence that appeared between 2013 and 2019. To our knowledge, no relevant systematic review has been published after 2013. The result, again, verified the clinical benefits of using a combination of SABA and IB in children and adolescents with asthma compared with using SABA alone and provided an evidence-base for clinical practice. However, this systematic review has several limitations. Firstly, because of the different diagnostic criteria of childhood asthma, the external validity of the studies is quite poor. Secondly, original research protocols were not always well described, which resulted in low quality of evidence. As such, there were some missing details for certain studies, which limited our ability to interpret the data. Thirdly, the applicability of results from the present review should be concluded with caution. The analyses of patients’ age, severity of asthma, and co-interventions were conducted with subgroup data from original studies and resulted consistent conclusion with 2013 review^14^. In addition, because of insufficient data, we were unable to perform subgroup analyses of other factors of interest, such as dosage regimens and frequency. Moreover, the treatment durations across the included studies varied. The differences may also affect the applicability of the present review results. Although data extrapolation from the non-randomized subgroup population should be cautious, the current conclusion of our meta-analysis may provide new ideas and directions to identify the clinical beneficiaries of combination therapy of IB + salbutamol. Further well-conducted and adequately powered RCTs with standardized outcome measures are needed to explore the most appropriate treatment dosage and duration for children and adolescents with asthma.

In conclusion, the results indicate that IB + salbutamol can significantly reduce the risk of hospital admission in children and adolescents, and this combined therapy may have significant clinical benefits in children with severe and moderate-to-severe asthma exacerbation. Future prospective randomized controlled trials are needed to evaluate the clinical benefits of combining IB and salbutamol in asthma children and adolescent in different age, severity of asthma, and co-interventions subgroups.

## Methods

### Registration

A prior protocol was developed and registered with PROSPERO (registration number: CRD42020159999). This review was informed by and reported using the Preferred Reporting Items for Systematic Reviews and Meta-Analyses guidelines ^16–18^ (E-Appendix 1).

### Search strategy

An electronic search for studies published up to 02/09/2019 using the key words “asthma,” “salbutamol,” and “ipratropium bromide” was conducted using the following databases: MEDLINE, Embase, Cochrane Library, China Biological Medicine Database, Chinese National Knowledge Infrastructure, Chongqing VIP, and Wanfang Chinese language bibliographic database. The search strategy was independently developed by two investigators according to the following selection criteria. Any dispute was resolved by mutual consensus with a third investigator.

Eligible clinical studies were defined based on the following criteria: (1) randomized controlled trials (RCTs); (2) children and adolescents aged ≤18 years; studies with mixed age population (both children and adults) were excluded if more than 5% of the participants did not meet the eligible criteria of age; (3) physician-diagnosed asthma by any appropriate diagnostic criteria (including wheezing children for <1 year); (4) comparing IB + salbutamol (either in a fixed dose or delivered separately) with salbutamol alone, without limitation of treatment duration, mode or frequency of administration, or dosage. Trials with additional treatments that were equally applied to both intervention and comparison groups were also included. There was no limitation of language.

### Outcome measures

The primary outcomes that were measured were hospital admission (as defined by original studies) and any adverse events. Secondary outcomes included pulmonary function (including percentage change from baseline of predicted % forced expiratory volume in one second [FEV_1_]; percentage change from baseline of FEV_1_; and change from baseline in respiratory resistance); clinical score (as defined by original studies, including accessory muscle, asthma, cough, dyspnea, wheeze, wheezing sound, and daytime/night-time symptoms scores); oxygen saturation; need for extra medication (including systemic corticosteroids and repeated bronchodilators); specific adverse events (including dry mouth, nausea, vomiting, and tremor); and relapse rate (defined as a return visit within a certain time that was predefined by original studies).

### Data extraction and assessment of risk of bias

Data extraction was independently performed by two reviewers. Missing data were requested from the corresponding authors via email for full original data. Any ambiguities in the selection and extraction were resolved by discussion, with the assistance from a third party if necessary. Once extraction was completed, data were reviewed to identify duplicate studies and duplicate reporting of populations and only the longest follow-up studies were retained. The extracted data included general study characteristics (including first authors, publication years, study center, and sample size); demographic characteristics (including diagnosis, age, and settings); interventions and controls (including frequency and treatment duration); and outcome characteristics (including categories and definitions of outcome and follow-up).

The Cochrane Risk of Bias tool ^18^ was applied to assess the quality of the included RCTs, including sequence generation, allocation concealment, blinding of participants and personnel, blinding of outcome assessment, incomplete outcome data, selective outcome reporting, and other potential threats to validity. Studies were rated on each variable as low risk, high risk, or unclear risk of bias. If a study received “high risk” judgment in any one domain, it would be classified as “high risk of bias”. Two independent assessors conducted quality assessment, and any disagreement was settled by reaching a consensus or consulting a third researcher.

### Data analysis

Data were synthesized and analyzed using RevMan version 5.3 ^16^. A random effects model was used to calculate pooled effect estimates comparing the outcomes between the intervention and control groups where feasible. Dichotomous outcome results were expressed as risk ratio (RR) with 95% confidence intervals (CI). Continuous scales of measurement were expressed as a mean difference. Heterogeneity was calculated using the I^2^ statistic. For I^2^≥50%, the heterogeneity was interpreted. Subgroup analysis was performed for primary outcomes based on the following characteristics: 1) age (<6, ≥6, 6–14, or >14 years); 2) severity of disease (mild, moderate, severe, mild-to-moderate, or moderate-to-severe, as defined by original studies); and 3) presence of combined co-intervention (with glucocorticoid, without glucocorticoid, with steroid, or with standard care, as defined by original studies) (for details, refer to E-Appendix 2). Sensitivity analysis was performed according to the description in protocol. Publication bias was analyzed using funnel plots for outcomes when there were more than 10 studies. Unfortunately, owing to inconsistent reporting of outcomes in pulmonary function and clinical scores, a meta-analysis could not be performed. Therefore, a descriptive synthesis of aforementioned outcomes was performed instead.

## Acknowledgements

The authors would like to thank Canghong Zhi and Dandan Zhao for providing medical writing support.

## Compliance with ethical standards

## Human Participants and/or Animals

This article does not contain any studies with human participants or animals performed by any of the authors.

## Informed consent

Not applicable.

## Supporting information

Supporting material is available online

---

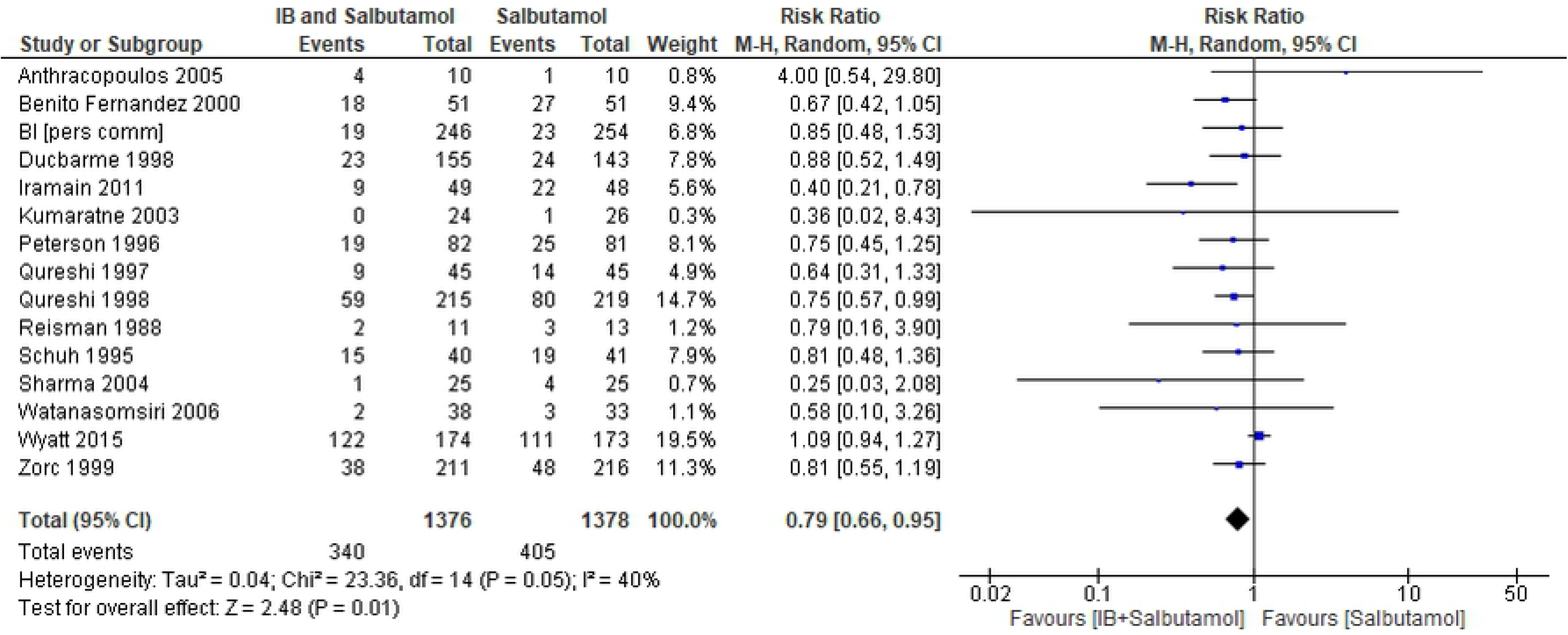

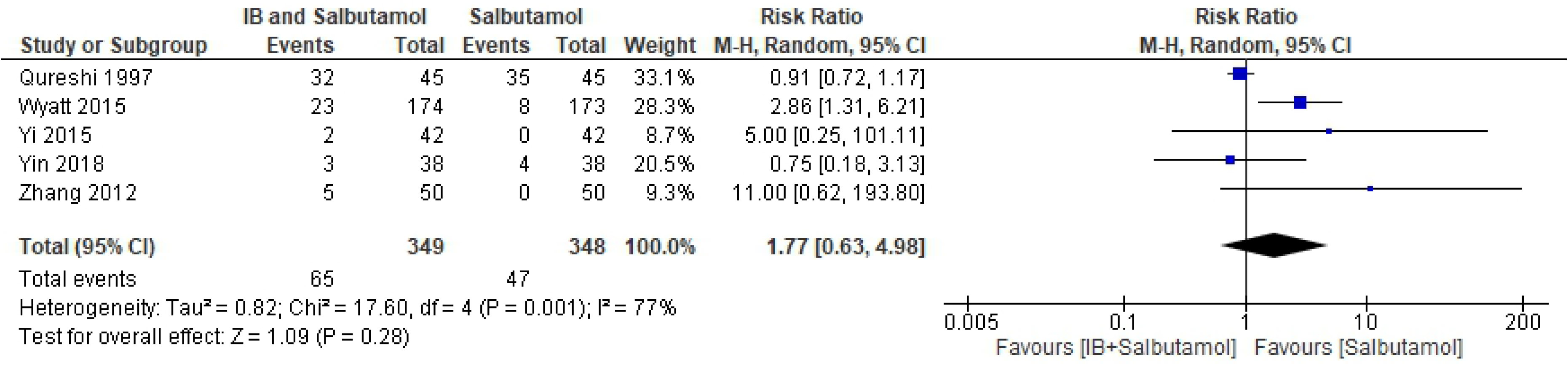

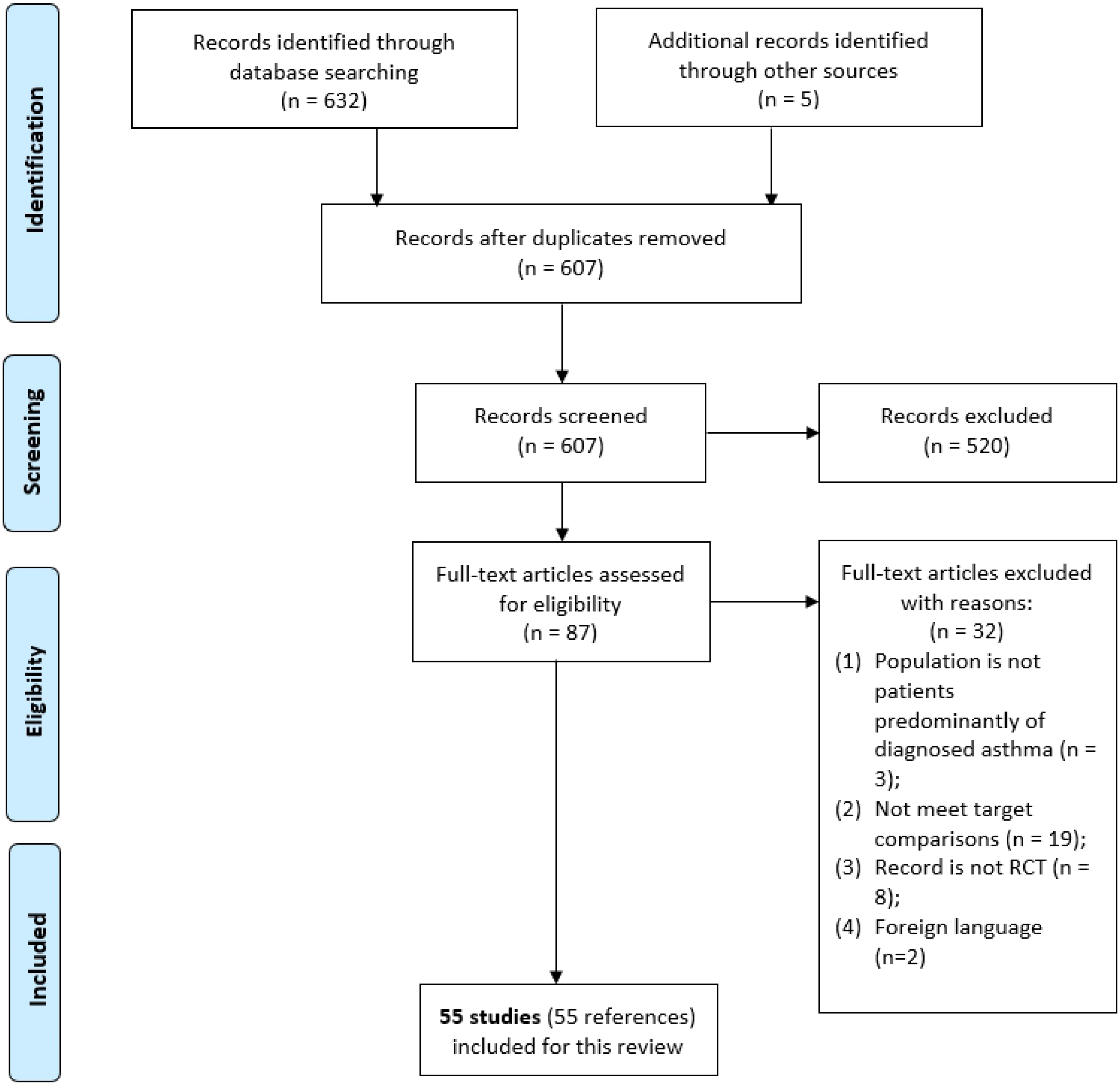

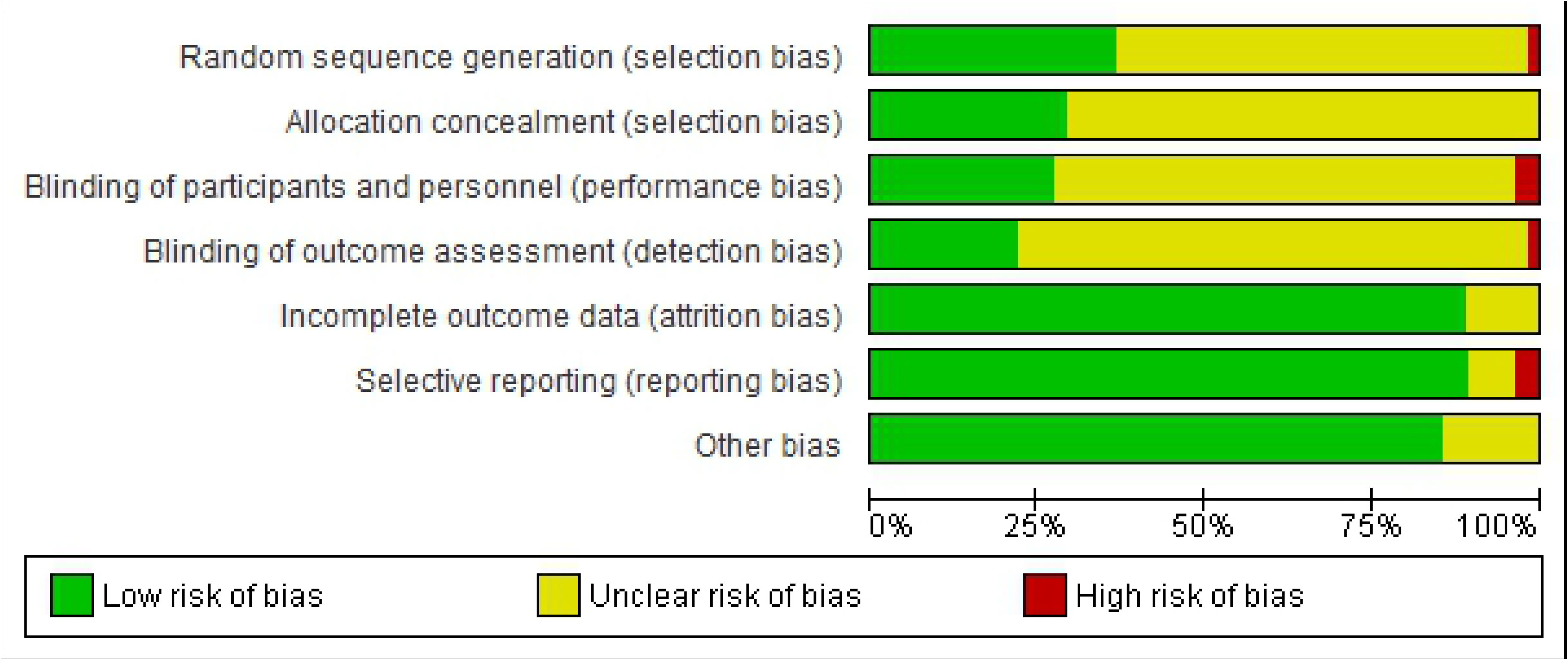

